# Advancing Target Discovery Through Disease-Specific Integration of Multi-Modal Target Identification Models and Comprehensive Target Benchmarking System

**DOI:** 10.1101/2025.08.06.668866

**Authors:** Howell Leung, Chengchen Duan, Wenhao Gou, Jianjiu Chen, Ying Xin, Zetian Zheng, Vladimir Naumov, David Gennert, Man Zhang, Alex Aliper, Feng Ren, Evgeny Izumchenko, Frank W. Pun, Alex Zhavoronkov

## Abstract

Target identification is crucial for drug development. AI-driven approaches leveraging multi-omics and computational modeling can accelerate this process. However, integrating multi-modal data for disease-specific target identification and predicting translational potential remains challenging. Moreover, the absence of a systematic evaluation framework for model performance limits confidence in target reliability. We present a unified platform combining machine learning-based target identification with comprehensive benchmarking. As a testbed, we developed Target Identification Pro (TargetPro), a disease-specific model spanning 38 diseases across oncology, metabolic, immune, fibrotic, and neurological categories. TargetPro shows strong predictive performance for clinical-stage targets and reveals disease-specific patterns, underscoring the need for tailored target detection models. We next created Target Identification Benchmark (TargetBench 1.0) to rigorously assess target identification systems, including large language models, based on their ability to recover established targets and find high-quality novel candidates. This integrated approach offers a streamlined strategy to evaluate target discovery models, ultimately improving drug development efficiency.

## Introduction

Drug development is hindered by high costs, inherent challenges, and a striking failure rate, as up to 90% of candidates entering clinical trials do not achieve regulatory approval^1–3^. Many of these failures can be traced to the earliest stages of drug discovery, particularly the selection of biological targets that later prove to be less effective or more toxic than anticipated^4^. Given that the total cost of developing a single new drug may reach several billion US dollars, early identification of the most promising targets and de-risking their development pathway are critical for the pharmaceutical industry and, ultimately, for public health^4^.

A notable effort for optimizing target identification is AstraZeneca’s five ‘R’s framework, comprising the right target, right patient, right tissue, right safety, and right commercial potential^5^. Introduced after an analysis of drug development projects conducted between 2005 and 2010, this framework places the ‘right target’ as its cornerstone, emphasizing targets with strong mechanistic links to disease and supported by robust biological and genetic evidence. Implementation of the 5R framework increased the success rate of early development (from candidate drug nomination to phase III trial completion) from 4% in 2005-2010 to 19% in 2012-2016^6^.

In parallel, the emergence of artificial intelligence (AI)-driven platforms has revolutionized target identification. Traditional methods, often relying on expert experience and limited datasets, suffer from inefficiencies and high attrition rates^7^. The AI-assisted target discovery approach uses machine learning and bioinformatics to integrate and analyze vast datasets encompassing genomics, proteomics, and literature, providing a systematic, evidence-based approach to target prioritisation ^8^. Examples of AI-driven target identification platforms include ConVERGE by Verge Genomics (See Related Links), TargetMATCH by Owkin (See Related Links), Open Targets by the European Bioinformatics Institute, the Wellcome Sanger Institute, and GlaxoSmithKline^9–11^, and PandaOmics by Insilico Medicine^12^. Notably, several targets prioritized by PandaOmics, which integrates multiple omics and text scores, have been validated in both preclinical and clinical studies^13–16^. These platforms underscore AI’s ability to bridge data-driven hypothesis generation and clinical translation, enhancing decision-making in drug discovery. Additionally, Large Language Models (LLMs) are being increasingly used in target selection. By analyzing vast amounts of scientific literature, LLMs can uncover hidden connections between genes, diseases, and targets. For example, BioGPT has been applied to discover potential dual-purpose targets that are relevant to both aging and age-related diseases, identifying novel anti-aging targets^17^. ChatGPT is also being explored for predicting protein domains, uncovering drug-binding pockets, and assisting in structure prediction^18^.

Despite these advances, two fundamental gaps continue to limit the full potential of AI in drug discovery. The first relates to the challenges of using multi-model platforms for disease-specific target identification and objective-driven prediction. Given the heterogeneous mechanisms underlying different diseases, it is unrealistic to expect a single model to provide a one-size-fits-all strategy for target identification across all disease contexts. However, for end users, selecting the optimal combination of target identification models is difficult due to the complexities of the underlying data sources and opaque assumptions inherent in AI-based approaches. This highlights the need for a validated, off-the-shelf framework for selecting and weighting multiple models tailored to specific disease contexts. In addition, current target identification systems lack the ability to predict a target’s potential to advance to clinical development, an objective that is important to end users. Developing machine learning models that can generate such predictions could assist human experts in making high-stakes decisions about which targets to prioritize for further development.

The second critical gap is the lack of standardized evaluation methods for these predictive tools. The rapid and heterogeneous proliferation of AI platforms has outpaced the development of common systems to evaluate their performance and reliability, especially for a specific task area. This ‘benchmarking gap’ slows scientific progress and industrial adoption, as it becomes difficult to compare different models, understand their relative strengths, or build the necessary confidence to integrate their outputs into multi-billion-dollar R&D decisions. Existing benchmarks, such as BETA, have been developed to address specific computational tasks like drug-target interaction prediction, but they may not fully capture the multifaceted biological and clinical considerations essential for selecting a viable therapeutic target for development^19^. Above all, a dependable and transparent evaluation system is essential for building trust and ensuring the responsible integration of AI into modern medicine.

This study introduces a framework designed to address these two gaps by integrating a machine learning workflow, PandaOmics Target Identification Pro (TargetPro), along with a dedicated evaluation system, Target Identification Benchmark (TargetBench 1.0). Together, these components form a strategy to improve the accuracy of target discovery and validate the reliability of target predictions (**Figure 1**).

**Figure 1.**
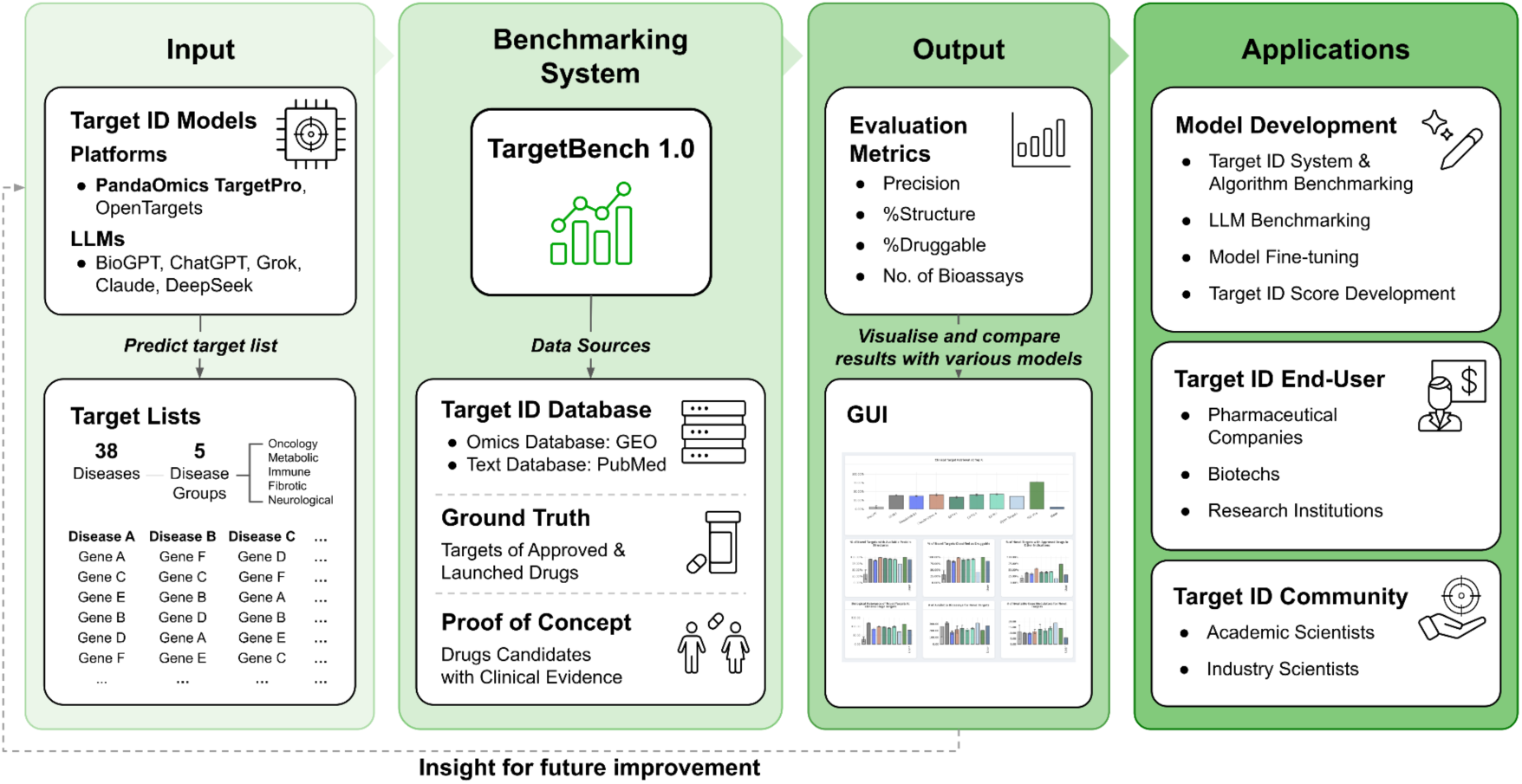
Overview of TargetBench 1.0 for drug target identification models benchmarking.

First, we developed TargetPro by implementing a disease-specific, objective-driven machine learning workflow designed to effectively utilize the multidimensional omics and text data derived from the PandaOmics platform^12^. TargetPro was trained using a meticulously curated set of targets that have entered the clinical stages (defined as Phase 1, Phase 2, Phase 3, and Launched) for a given disease. By using these targets as training labels, TargetPro learns the complex patterns and features associated with successful translation, enabling it to predict targets with a higher probability of advancing through clinical development. Second, we established TargetBench 1.0, a system designed to systematically evaluate the performance of any target discovery model and platform. This benchmark aligns with the practical goal of identifying targets with the potential to advance to therapeutic application, providing a clear and relevant measure of predictive power.

We applied this framework to 38 representative diseases, selected based on their prevalence, incidence, and data availability, spanning oncology^20^, immune-related^21^, metabolic^22^, fibrotic-related^23^, and neurological disorders^24^. TargetPro generated 38 disease-specific target lists, prioritizing those with the highest potential for clinical advancement. These lists were then compared against outputs from other target identification models (including LLMs) using TargetBench 1.0, which provides a comprehensive assessment across multiple dimensions such as: clinical target retrievability, structural data availability, druggability potential, repurposing opportunities, biological relevance, bioassay accessibility, and gene modulator availability. Collectively, this integrated approach enhances the accuracy of target discovery, establishes confidence in computational predictions, and ultimately accelerates the development of novel therapies.

## Materials and Methods

### Data collection and preprocessing

Representative disease selection

We selected a comprehensive set of 38 diseases to represent a broad range of pathological mechanisms and therapeutic areas, guided by their prevalence, incidence, and data availability. These diseases were manually curated and systematically categorized into five major pathophysiological categories: oncology, metabolic disorder, immune-related disease, fibrotic-related disease, and neurological disease. This diverse selection enables robust and generalizable comparison of drug target characteristics across distinct biological contexts. The specific diseases included in each category are summarized in **Supplementary Table 1**. The oncology group comprised 18 malignancies: non-small cell lung carcinoma, breast cancer, prostate cancer, colorectal cancer, thyroid cancer, gastrointestinal cancer, bladder cancer, endometrial cancer, cervical cancer, acute leukemia, melanoma, renal cancer, liver cancer, ovarian cancer, myeloma, pancreatic cancer, chronic myelogenous leukemia, and head and neck cancer. The metabolic disorder group included 5 diseases: obesity, hypertension, type 2 diabetes mellitus, hyperlipidemia, and chronic kidney disease. The immune-related category consisted of 6 disorders: inflammatory bowel disease, rheumatoid arthritis, systemic lupus erythematosus, psoriasis, type I diabetes mellitus, and scleroderma. The fibrotic-related group contained 3 pathologies: idiopathic pulmonary fibrosis, atherosclerosis, and osteoarthritis. Finally, the neurological disease category included 6 conditions: Alzheimer’s disease, Parkinson’s disease, amyotrophic lateral sclerosis, stroke, epilepsy, and Huntington’s disease.

Clinical-stage targets annotation

For each disease, the analysis encompassed 19,291 protein-coding genes annotated in the HGNC database (See Related Links). A manually curated disease-drug-target list was compiled from the latest public sources (e.g., ClinicalTrials.gov) and was subsequently reviewed and verified by our internal team of biologists. This manually curated list was used to assign each target to a specific stage of drug development (Preclinical, Phase 1, Phase 2, Phase 3, and Launched). Preclinical targets are defined as all protein-coding genes, excluding those in Phase 1-3 clinical trials or currently launched. Drugs with unspecified targets or without a clear mechanism of action were excluded. If multiple drugs targeted the same protein, the protein was assigned the highest development stage among its associated drugs.

Target identification model scores

We collected 22 target identification model scores for 38 disease indications from PandaOmics, an AI-driven target discovery platform frequently used in target identification studies^12–14,25^. The scoring system includes 12 omic-based metrics (expression, pathways, causal inference, network neighbors, interactome community, knockouts, disease submodules, overexpression, matrix factorization, heterogeneous graph walk, mutated submodules, and mutations) and 10 text-driven scores (attention score, credible attention index, grant funding, mean hirsch, impact factor, evidence, grant size, attention spike, funding per publication, and trend). The omics and text scores are derived from sophisticated algorithms that integrate and analyze data from genomics, transcriptomics, proteomics, publications, patents, grants, and clinical trials^12^.

### Machine learning

#### Model training for TargetPro

We developed target identification models (TargetPro) using XGBoost classifiers^26^ within a positive-unlabeled learning framework^27^. This approach was chosen to address the inherent challenge in drug discovery, where definitive labels (i.e., targets confirmed to be irrelevant to a disease) are often unattainable. In our PU-learning setup, we defined targets with clinically developed drugs from commercial entities as the ‘positive’ class. All other targets (also known as unlabeled targets), which could be true negatives or currently undiscovered positives, were treated as ‘negative’.

To address the significant class imbalance between the ‘positive’ and ‘negative’ sets, we applied class weights inversely proportional to their frequencies:

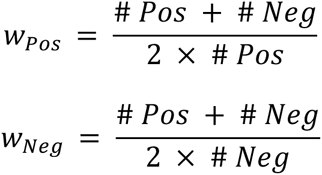

Where *w*_*Pos*_ and *w*_*Ne*g_ are weights to positive and negative class; # *Pos* and # *Neg* are numbers of observations in positive and negative classes, respectively.

A distinct model was trained for each indication, recognizing that different biological patterns govern target success in different therapeutic areas. Model training and hyperparameter tuning for each of these disease-specific models were conducted using a nested 5-fold cross-validation approach (**Figure 2A**). The entire dataset was first partitioned into five equal folds. The main process was then repeated five times, with each iteration using four folds for model training and holding out the remaining fold as a final, unseen test set. Within each training repetition, a separate inner cross-validation loop was performed exclusively on the training data for hyperparameter tuning. The optimal hyperparameters were subsequently used to train a single model on the entire training set, which then made predictions on the held-out test fold. Finally, the out-of-sample predictions from all five folds were combined to assess the model’s overall performance using the Area Under the Precision-Recall Curve (AUPRC), a metric that is particularly well-suited for evaluating classifiers on imbalanced datasets.

**Figure 2.**
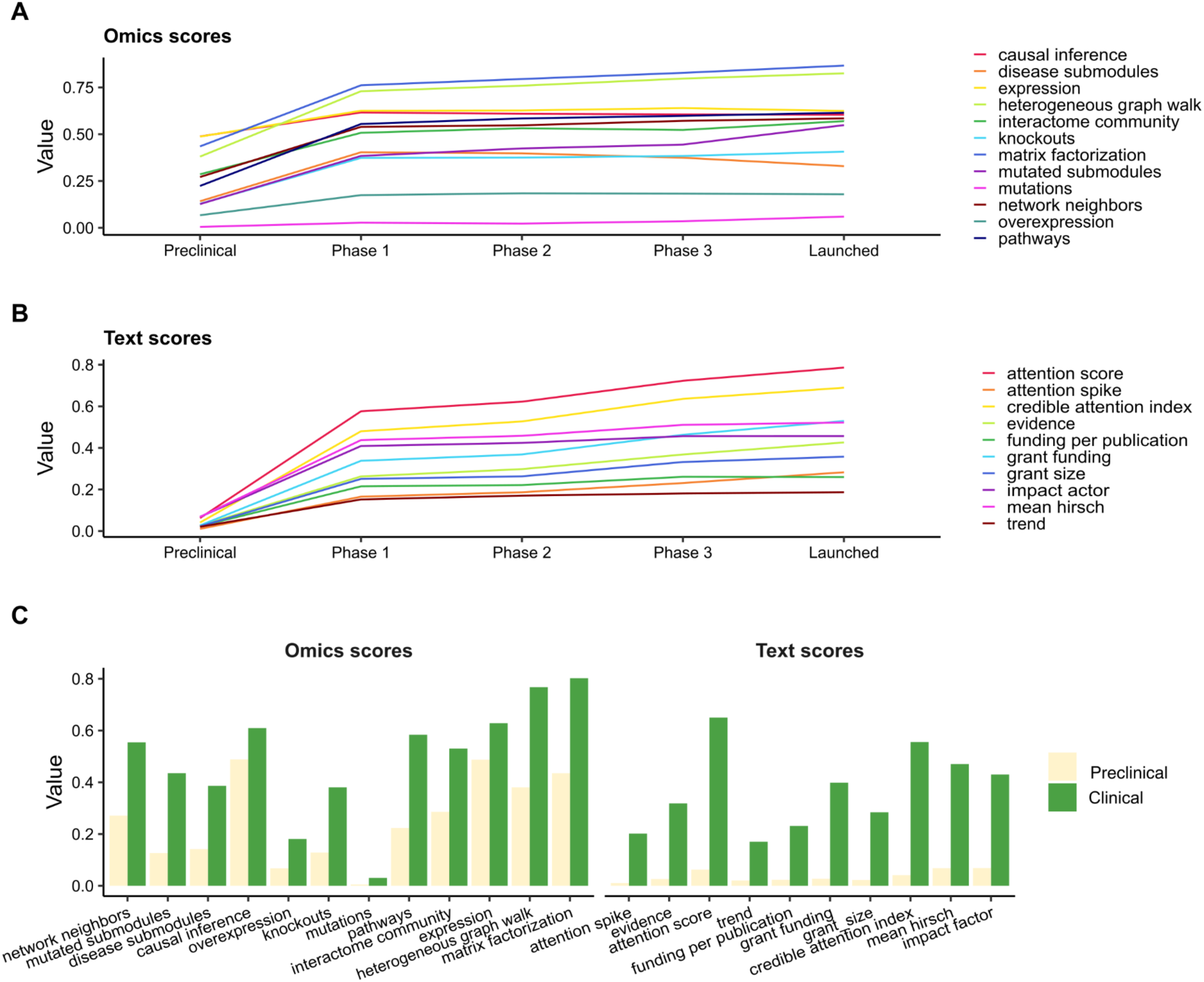
Omics and text scores across clinical development stages. All plots are generated using the combined scores from all 38 diseases to provide a global overview. (A, B) Trend plots of the 22 omics and text scores across distinct stages of clinical development, from ‘preclinical’ to ‘launched’. (C) Bar plots comparing the mean of scores for targets grouped as ‘preclinical’ versus ‘clinical’ (Phase 1 and beyond). All models exhibit significant differences (Wilcoxon rank-sum test, P-values < 0.05).

#### Feature importance analysis using SHAP framework

We implemented the Shapley Additive Explanations (SHAP) framework^28^ to provide an interpretable and consistent feature importance analysis. Global feature importance was determined by calculating the mean absolute SHAP value for each feature, representing the average magnitude of its contribution to the model’s predictions across the entire dataset. To facilitate a higher-level analysis, individual features were subsequently aggregated into three categories based on their data source and nature: Text, Omics (Static), and Omics (Dynamic). This allows the assessment of the relative influence of different data modalities on model performance. The Text category included features derived from publication and funding metadata, such as attention, credible attention index, grant funding, mean hirsch, impact factor, evidence, grant size, attention spike, funding per publication, and trend. The Omics (Static) category comprises features whose values are calculated independently of the golden and user’s proprietary datasets. These features, representing stable biological states and pre-computed network properties, included matrix factorization, heterogeneous graph walk, mutated submodules, and mutations. In contrast, the Omics (Dynamic) category consists of features whose scores are directly affected by the specific input of omics data for an analysis. These features capture dynamic or latent biological signals and include expression, pathways, causal inference, network neighbors, interactome community, knockouts, overexpression, and disease submodules.

### Benchmarking

#### Setup for model benchmarking

To benchmark different target identification models, we conducted a comparative evaluation of the therapeutic target lists generated by each model. To ensure a fair and meaningful comparison, the experimental setup for each model was tailored to mimic its typical real-world application and maximize its performance. For the LLM evaluation, a standardized prompt (see Prompt for querying LLMs) was used to query each model (BioGPT, Grok3, DeepSeek-R1, Claude-Opus-4, GPT4o, GPT4.1, GPTo4 mini, and GPTo3) for a list of targets for each of the 38 diseases. To account for the stochastic nature of these models, the process was repeated five times per disease. We prompted LLM to nominate a defined number of genes per iteration and then aggregated results across all runs for the final analysis. For Open Targets, we retrieved disease-specific target lists utilizing their default ranking, which integrates genetic, omics, and text-based evidence. The evaluation protocol for TargetPro was specifically designed to simulate a user performing a new meta-analysis by integrating additional, user-selected datasets with the foundational ‘golden dataset’. For each disease, we augmented the baseline ‘golden dataset’ with additional disease-specific omics data, creating an enhanced ‘golden-plus dataset’. TargetPro then generated new target prediction scores using the combined dataset. See **Supplementary Figure 1** for details of the prediction process.

#### Prompt for generating therapeutic targets

To elicit a ranked list of potential therapeutic targets for each disease, the following standardized prompt was provided to each LLM, with {disease_name} and {top_k} parameters adjusted for each query:

’For {disease_name}, generate a complete list of {top_k} high-confidence drug targets, where each target is a gene name. High-confidence targets are defined as: Genes with drugs in clinical-stage development (e.g., approved or in trials) for {disease_name}, AND/OR preclinical targets supported by substantial evidence of their role in {disease_name} and their potential as drug targets. You should rank these {top_k} targets by confidence level.

Your response should be a json file ONLY. The JSON file should have {top_k} elements only, with each element only storing 1 gene name, ordered by their rank. Ignore the alternative name of each gene in your output. Ignore any unrelated content.’

#### Assessment of robust target identification models

The robustness of the target identification models is evaluated based on their ability to enrich top-ranked predictions with high-quality attributes. In real-world applications, from the perspective of target biologists and drug hunters, confidence in a model’s output is initially established by the presence of well-validated targets^29^ . Consequently, the primary metric for assessing model quality is the clinical target retrieval at top K, which can be expressed with the following equation:

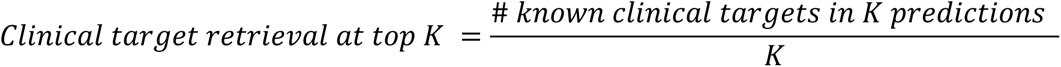

This quantifies the retrieval of established targets within the top-ranked predictions and serves as a crucial benchmark for the model’s veracity, where K is the number of established clinical targets for the disease.

However, we acknowledge the inherent paradox in drug discovery: while retrieving known targets validates a model’s accuracy, the discovery of promising novel targets often holds greater translational value. Therefore, a comprehensive evaluation must be extended to assess the intrinsic quality of all identified targets, whether known or novel. This ensures that the model is not only accurate but also capable of generating candidates with a high probability of success in the development pipeline.

Beyond the clinical target retrieval, we also evaluated the quality of each model’s novel predictions. We focused on novel targets, defined as those ranked within top K but not yet in clinical development, as this reflects the practical need in reality to prioritize a limited number of high-potential candidates. These novel targets were assessed using a suite of metrics designed to quantify key characteristics relevant for drug development. We established a multi-faceted set of criteria, sourced from expert-curated databases, to ensure the robustness and clinical relevance of the identified targets. These criteria assess targets across several critical dimensions, including their clinical development stage, scientific validation, and safety profiles derived from clinical trial data and biological essentiality assessments.

First, the availability of protein crystal structures was confirmed through the Protein Data Bank, as this information is crucial for facilitating subsequent drug design to improve potency and selectivity. Next, a target’s druggability was systematically evaluated using information from multiple resources (Citeline, DGIdb & DrugBank). Gene targets with drug records are considered druggable, prioritizing those with a high likelihood of successful pharmacological modulation to ensure chemical and biological feasibility. To assess therapeutic adaptability and de-risk development, we identified targets with approved drugs in other indications by referencing comprehensive drug development and clinical trial databases, leveraging their established safety and efficacy profiles. To connect targets to established disease-specific biological mechanisms, their biological relevance was measured by the number of pathways they share with existing clinical-stage targets, based on data from Reactome. The level of experimental validation for each potential target was quantified by the number of available bioassays cataloged in PubChem, with a higher number of associated assays indicating a more robust evidence base and greater development potential. Finally, we used MedChemExpress (See Related Links) to obtain the number of available gene modulators, which included small molecules, peptides, and monoclonal antibodies, but excluded siRNA.

### Use of AI in manuscript preparation

During the preparation of this manuscript, AI tools were utilized to assist with writing and editing. The role of these AI tools was limited to proofreading for grammatical errors, improving sentence structure, and enhancing overall clarity and readability. The authors carefully reviewed and revised all AI-generated suggestions to ensure scientific accuracy and retain full responsibility for the final content of this publication.

## Results

### Characteristics predictive features

We applied our target identification and benchmarking framework to a curated set of 38 diseases spanning five major therapeutic areas: oncology, metabolic, immune-related, fibrotic-related, and neurological diseases. The initial analysis focused on characterizing the foundational datasets for this diverse group. Among the 38 selected diseases, considerable heterogeneity was observed in both the number of clinical-stage targets (used as positive labels in model training) and in the ratio of clinical-stage to preclinical targets (used as negative labels and defined as all protein-coding genes excluding those in Phase 1-3 clinical trial or currently launched) (**Supplementary Table 1**). The oncology group contained the largest number of clinical-stage targets (ranging from 92 to 453), whereas other groups, such as metabolic, had fewer (ranging from 71 to 180). The class imbalance between positive and negative labels also varied substantially, ranging from 1:42 in oncology to 1:356 in neurological diseases. These observations underscore the sparsity of validated targets among certain disease areas and the striking variability in target landscapes across different pathologies.

We next analyzed the behavior of 22 distinct feature scores, which quantify a target’s association with disease using evidence from omics data, scientific literature, clinical records, and grants (see **Materials & Methods** for details), across the different stages of clinical development. We observed that the magnitude of most scores increased progressively from the ‘preclinical’ stage into Phase 1, after which it remained relatively stable (**Figures 2A and B**). A clear distinction between ‘preclinical’ and ‘clinical’ stages emerges when targets are grouped accordingly (**Figure 2C**, Wilcoxon rank-sum test, P-values < 0.05), indicating that these scores provide predictive insights for the transition into clinical development.

### Workflow implementation and TargetPro development

Given the distinct feature patterns differentiating preclinical and clinical-stage targets, we hypothesized that a machine learning model could be trained to identify targets with a higher likelihood of advancing to the clinical stage. The model is based on the assumption that targets with a high predictive score (TargetPro score) share key characteristics with known clinical-stage targets and are therefore more likely to advance into clinical development. To construct this model, we implemented a systematic machine learning workflow that integrates 22 omics and text scores and uses developmental stage labels assigned to each gene as ground truth for supervised learning (**Figure 3A**; see **Materials and Methods** for full details).

**Figure 3.**
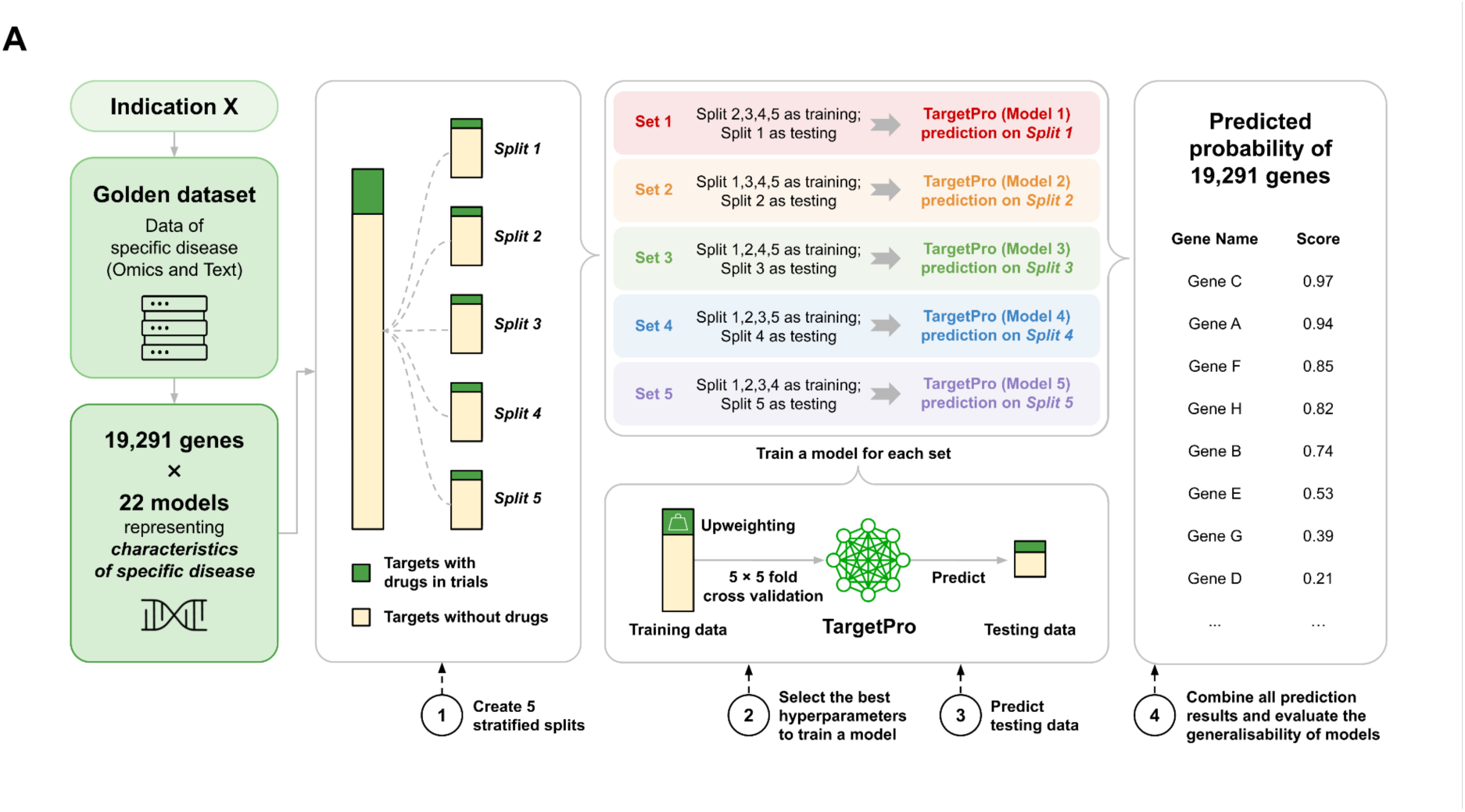
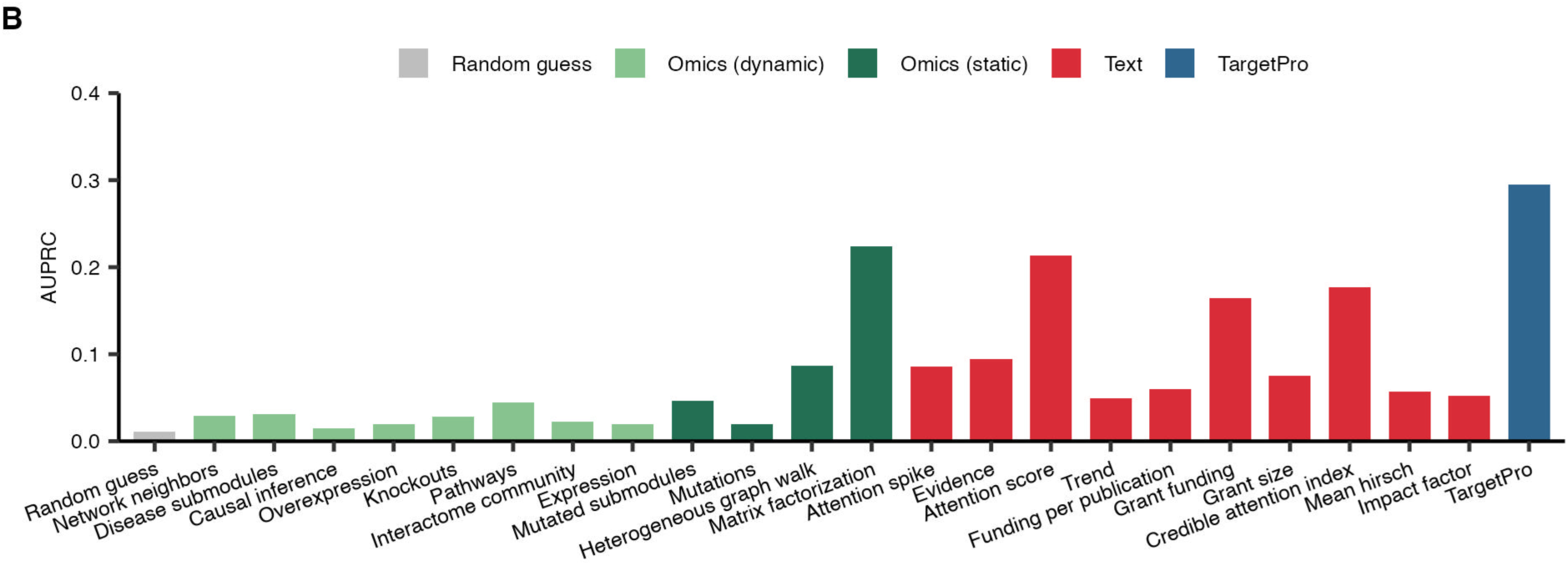
(A) Workflow for TargetPro model training. The process begins with a manually curated ‘Golden dataset’, representing the most relevant and available data for a given disease. This dataset is partitioned into five stratified splits based on whether targets have associated drugs in clinical trials. A nested 5-fold cross-validation is then employed, where an inner loop performs hyperparameter tuning with class upweighting. A TargetPro model is then trained with the optimal parameters to generate predictions on held-out testing data. Finally, the out-of-sample predictions from all five folds are combined to evaluate the overall generalizability of the model. (B) Performance comparison of the integrated TargetPro model against each of the 22 individual omics and text models.

With this foundation, we trained a distinct TargetPro model for each therapeutic area using an XGBoost algorithm^26^. To optimize performance and account for the inherent class imbalance between clinical-stage and preclinical targets, we integrated an upweighting strategy and conducted disease-specific hyperparameter tuning. This disease-specific design is a core feature of our workflow, recognizing that distinct biological patterns govern target progression in different therapeutic areas. The framework’s flexibility enables it to capture these nuances while maintaining rigorous and consistent methodological standards. Ensuring that the model produced interpretable weights for each input score allowed us to examine the features driving the predictions and compare their contributions across therapeutic areas.

To ensure a robust and unbiased assessment, model performance was evaluated using a nested 5-fold cross-validation strategy, yielding predictions for all genes. The Area Under the Precision-Recall Curve (AUPRC) for TargetPro was significantly higher than that of any single omics and text score used as a baseline (AUPRCs: 0.29 vs. 0.015-0.22, paired T-tests, P-values < 0.05, **Figure 3B**), reflecting superior predictive power in the clinically relevant precision-recall space.

### Disease-specific patterns of feature importance

To understand the predictive drivers of our models, we analyzed feature importance across the five disease groups (**Figure 4**). At the individual feature level, matrix factorization and attention score were universally the most impactful, highlighting their core contribution to the trained disease-specific TargetPro models.

**Figure 4.**
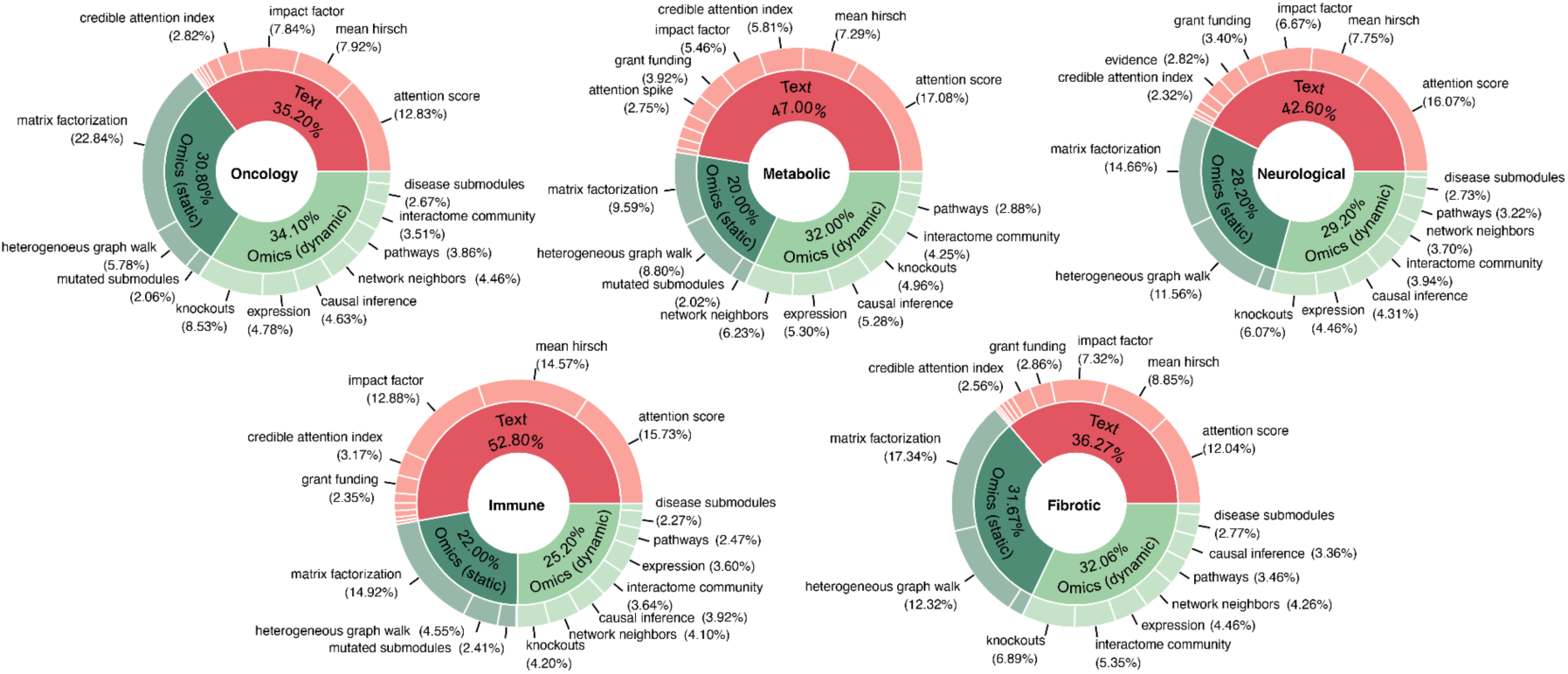
Features importance of TargetPro models across five disease groups, only features with importance ≥ 2% are labelled.

However, the relative importance of these top features varied by disease context. Matrix factorization was the single most important feature for the oncology (22.84%) and fibrotic (17.34%) models. In contrast, attention score was the leading contributor for the immune (15.73%), metabolic (17.08%), and neurological (16.07%) models. Other features also showed specific utility, such as heterogeneous graph walk in neurological (11.56%) and fibrotic (12.32%) models. Additionally, the knockout score demonstrated a clear pattern in its contribution, ranking highest in oncology (8.53%), followed in descending order by fibrotic (6.89%), neurological (6.07%), metabolic (4.96%), and immune models (4.20%).

When features were aggregated into higher-level categories, a clear interplay between data sources became apparent. Features derived from text were highly influential, representing the largest single category of importance in the immune (52.80%), metabolic (47.00%), and neurological (42.60%) models. The oncology and fibrotic models, however, demonstrated a more balanced dependency on all three feature categories, with nearly equal contributions from text, omics (Static), and omics (Dynamic) sources (see **Materials and Methods**). The contribution of omics-based features was substantial across all groups and accounted for the largest share of importance in several models. In particular, omics features collectively account for 64.90% and 63.73% of the total importance in the oncology and fibrotic models, respectively, surpassing the contribution from text-based features.

### Model performance assessment in target identification

To address the existing lack of standardized methodologies for evaluating target identification models, we developed TargetBench 1.0. This platform rigorously assesses the performance of various models in target prediction, simulating their real-world application environment (see **Materials and Methods**). The complete target lists for all 38 diseases across different models are provided in **Supplementary Tables 2 and 3**. TargetBench 1.0 is accessible at https://www.targetbench.org, enabling users to systematically benchmark their own target identification model using the metrics described below and demonstrated in **Supplementary Figure 2**.

We first evaluated the models’ ability to retrieve established clinical targets, which represent the most reliable ground truth and provide confidence to biologists and drug developers in real-world applications, by measuring the proportion of known clinical targets present among the top-ranked predictions from each platform (**Figure 5A**). TargetPro achieved an overall clinical target retrieval at top K of 71.6%. This corresponds to a 2-3 fold improvement over the tested LLMs, including GPT-4o and DeepSeek-R1, which achieved a retrieval rate of clinical targets in the 15-40% range. Our model also significantly outperformed Open Targets, which scored just under 20%. Notably, the strong performance of TargetPro was not confined to a single domain but remained consistently high across diverse therapeutic areas, including oncology, metabolic, immune, fibrotic, and neurological diseases.

**Figure 5.**
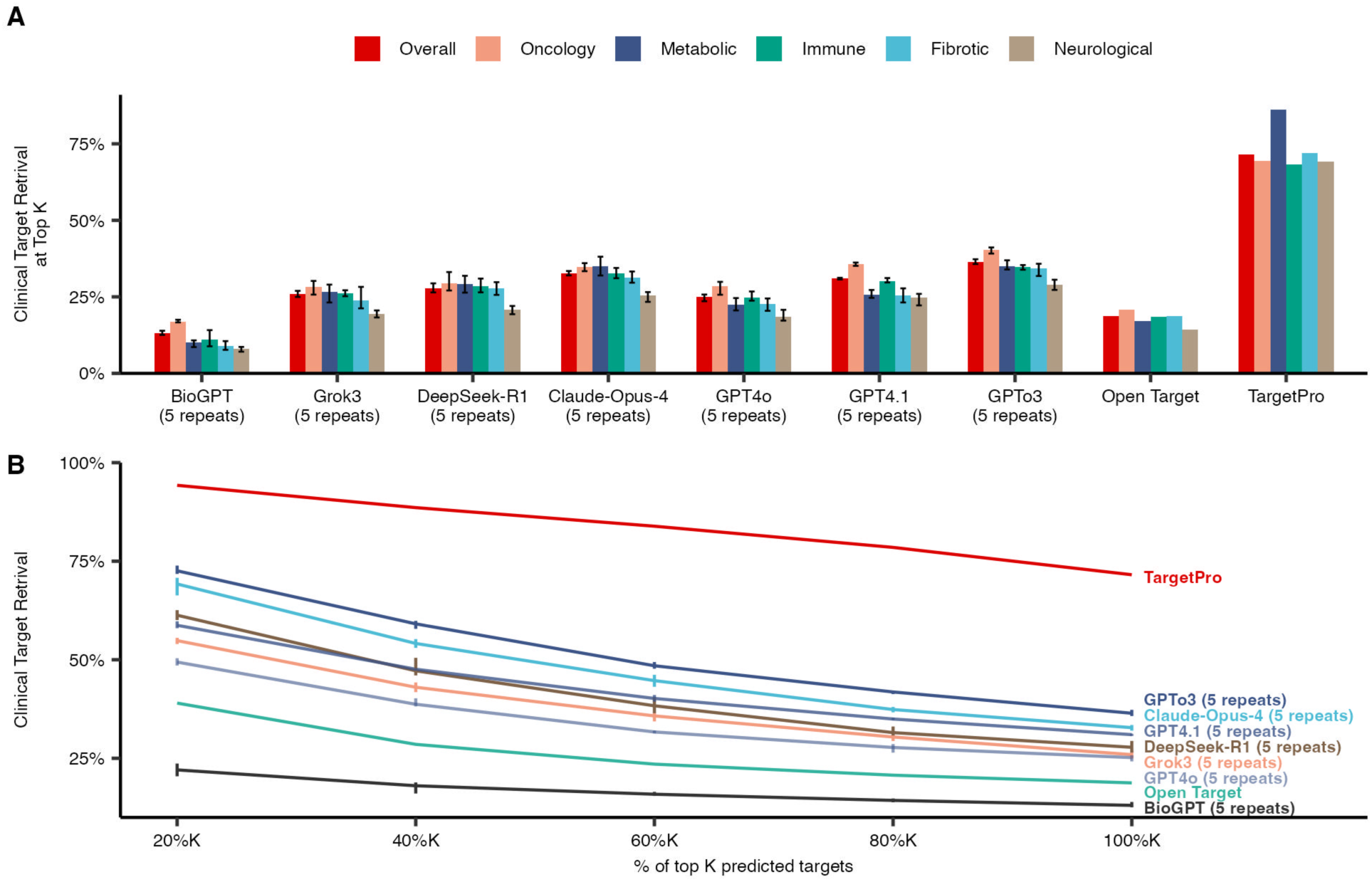
Model performance in clinical target retrieval at (A) top K Targets and (B) top K Percentage, where K is defined as the number of established clinical stage targets of a specific disease.

Next, we investigated how the number of requested targets, a critical factor from the user’s perspective, affects the clinical target retrieval performance (**Figure 5B**). TargetPro consistently demonstrated the highest performance across different percentages of K. Its precision showed a slight gradual decline starting at above 95% when users requested 20% of K targets, and remained as high as 71.6% when considering 100% K. In contrast, all other evaluated models, including the suite of LLMs and Open Targets, exhibited a much steeper decline in precision as the number of requested targets increased. For instance, some of the higher-performing LLMs (such as GPTo3 and Claude-Opus-4) started with a precision of nearly 70% but dropped by over 25 percentage points when 100% K was requested. Other models, such as GPT4o and Open Targets, started at more moderate precision levels (< 50%) and also showed a consistent decline, while BioGPT exhibited the lowest precision across all evaluated percentages of K.

While metrics such as clinical target retrieval at top K are useful for evaluating a model’s ability to recover known clinical targets, the real-world utility in drug discovery often lies in identifying promising novel candidates. We therefore focused our analysis on evaluating the characteristics that are critical for advancing novel targets through the early stages of the discovery pipeline. For this assessment, we examined the novel targets (defined as the top K predictors after excluding known clinical-stage targets) from each model and analyzed their druggability, biological relevance, and potential for drug repurposing.

First, in modern therapeutic development, having a target with a known structure is a crucial accelerator. We found that TargetPro excels here, with 95.7% of its proposed targets having an available 3D structure in the PDB, providing a clear advantage over other high-performing LLMs, which range from 60.3 to 91.3%. This advantage, combined with its stable and uniformly high performance across all therapeutic areas (91.7-99.6%), underscores TargetPro’s reliability in generating structurally enabled targets ready for drug design campaigns (**Figure 6A**). Additionally, a target’s druggability enhances the potential for therapeutic development, particularly for small-molecule drugs. TargetPro outperforms other LLMs by identifying a higher proportion of druggable targets with clinical evidence (86.5% vs 38.8-69.8%, **Figure 6B**). Furthermore, we evaluated the repurposing potential of novel targets, defined as the percentage of targets with approved drugs in other indications. TargetPro demonstrated a remarkable advantage, with 46% of its targets falling into this category, significantly outperforming all other platforms whose outputs were typically in the 17-28% range (**Figure 6C**).

**Figure 6.**
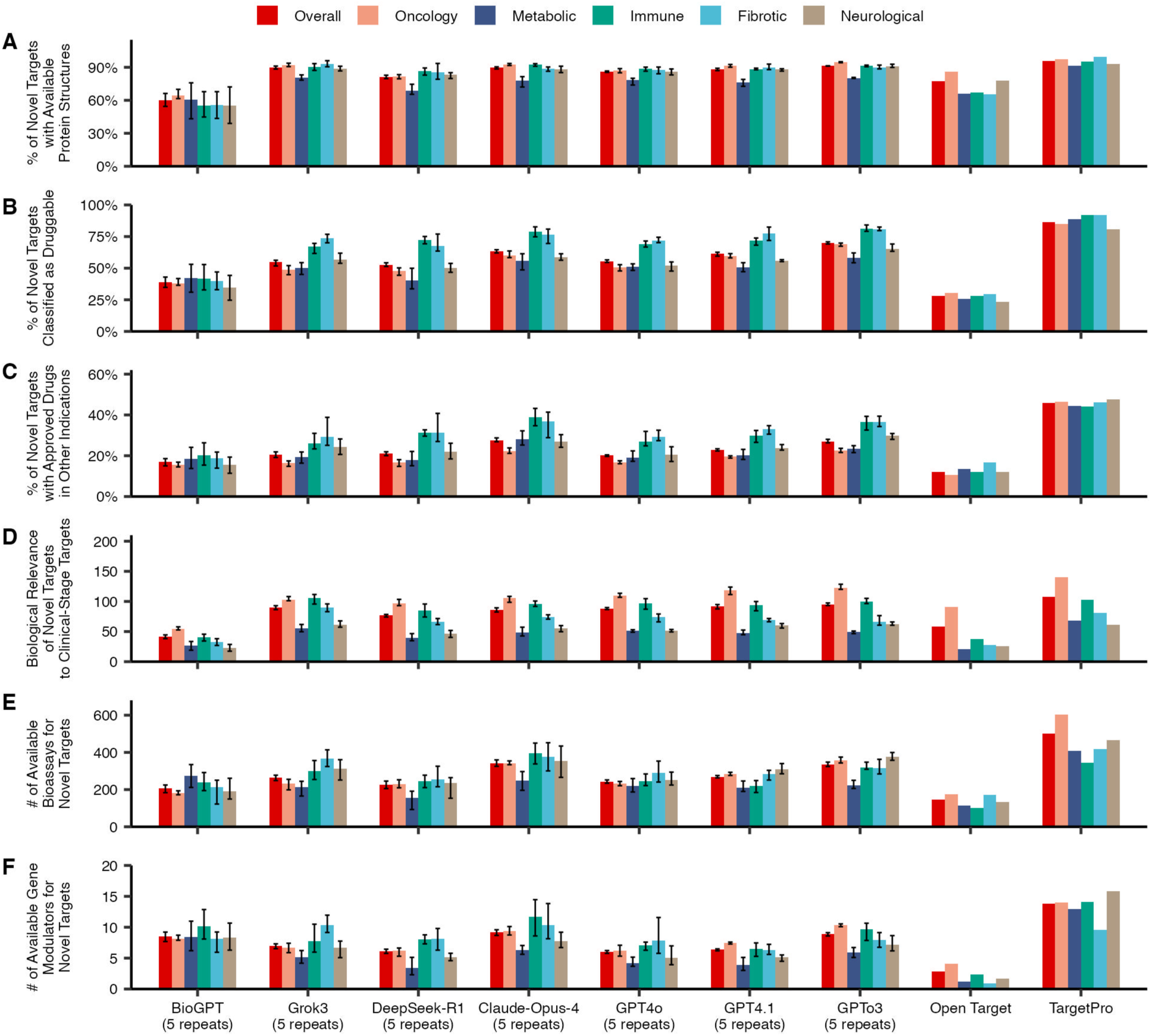
Multi-dimensional evaluation of novel targets selected by different target identification models: (A) the percentage of targets with available protein crystal structures; (B) the percentage of targets classified as druggable, supported by clinical evidence; (C) the percentage of targets with approved drugs in other indications; (D) the biological relevance, measured by pathway overlap with clinical-stage targets; (E) the average number of available bioassays; and (F) the average number of available gene modulators.

Beyond druggability, a target must be biologically relevant to the disease. We assessed this relevance by examining the extent of pathway overlap with established clinical-stage targets. In this analysis, TargetPro performed on par with the latest generation of LLMs, showing over 100 overlaps, while the overall pathway overlaps in Open Targets fell below 100 (**Figure 6D**). We also evaluated the feasibility of experimental validation. Targets identified by TargetPro had a vastly greater number of available bioassays (averaging over 500), a more than 1.4-fold increase compared to any other model (**Figure 6E**). Finally, the availability of gene modulators would facilitate the development of new drugs. Targets prioritized by TargetPro had more modulators on average than targets prioritized by LLMs (14 vs 6-9, **Figure 6F**).

## Discussion

AI-driven target identification platforms and LLMs have been gradually adopted to enhance the efficiency of target discovery. However, two long-standing challenges remain. First, applying multiple-model strategies for disease-specific target identification is difficult, as there is no established framework for selecting or optimally integrating these models. Secondly, there is a lack of transparent and systematic approaches for evaluating the performance of target identification platforms. This study puts forward a comprehensive framework designed to address these fundamental challenges. Specifically, our proposed solution consists of two main components: TargetPro, which automatically determines optimal feature combinations to enhance target prediction, and TargetBench 1.0 system, which comprehensively evaluates model’s predictive performance.

This integrated approach is designed to generate what can be termed actionable intelligence: reliable, well-characterized, and prioritized target hypotheses that can guide subsequent experimental work^30^. The central premise is that predictive accuracy and the ability to retrieve evaluative clinical targets are not separate objectives but rather complementary aspects of the same process. By developing these components together (**Figure 1**), we establish a positive feedback loop where better models drive the need for more sophisticated benchmarks. In turn, these benchmarks provide the confidence required to deploy the models in real-world drug discovery pipelines, ultimately aiming to reduce the high failure rates that currently affect drug development.

The design of the TargetPro component is guided by a detailed analysis of the characteristics and limitations underlying the data. This analysis confirmed that the data describing individual targets contain valid signals of clinical potential. Specifically, there is a positive association between each model and a target’s progression through clinical development, with all 22 features showing a statistically significant difference between preclinical and clinical-stage targets (**Figure 2**). This finding is consistent with the established understanding that most genetic associations and pathway analyses for preclinical targets remain poorly characterized (supported by limited evidence), whereas clinical-stage targets are extensively studied and have well-defined mechanisms of action (MOA)^31^. However, the utility of these individual scores for predicting success between clinical phases appears limited, as most scores tend to plateau once a target enters clinical stages. This plateau likely reflects the fact that subsequent progression depends not only on the target’s intrinsic biology but is also heavily influenced by external factors such as compound efficacy, trial design, human-specific safety profiles, regulatory considerations, and even commercial priorities^4^.

We therefore framed the problem as a binary classification task, distinguishing preclinical from clinical-stage targets, and trained machine learning models predicting a target’s overall potential entering the clinical development pipeline. The effectiveness of this strategy is demonstrated by the model’s performance. The trained TargetPro models achieved an AUPRC of approximately 0.3, significantly outperforming any individual omics and text scores (**Figure 3B**). These results indicate that a well-trained machine learning model can effectively integrate individually weak signals to enhance drug target discovery. This aligns with a growing body of evidence showing that multi-omics data integration can reveal complex biological patterns that are not detectable from any single data type alone^32^. In addition to improving predictive performance, evaluating the contribution of different features provides insight into the mechanisms underlying target prioritization. The SHAP analysis presented in **Figure 4** reveals that TargetPro’s decision-making is nuanced and context-dependent, with feature importance varying across disease groups. These results indicate that the model does not rely on simple, fixed rules but instead learns biologically relevant, disease-specific patterns. For instance, the contributions of omics scores vary across disease types, with the strongest contributions observed in oncology, followed by fibrotic, neurological, and metabolic diseases. In contrast, immune-related disorders show a weaker impact from omics data, likely reflecting the systemic and highly adaptive nature of immune responses. These findings align with established biological hallmarks of each disease group. Cancer is characterized by genome instability, frequent mutations, and widespread transcriptional dysregulation, making omics information particularly valuable for discovering anti-cancer targets^33^. Conversely, the complex and dynamic interplay between immune cells, cytokines, and environmental factors may limit the predictive utility of omic scores in immune-related diseases^34^.

Furthermore, several features consistently rank highly in importance across all disease groups. These include: (i) matrix factorization (9.59-22.84%), a metric for hidden gene–disease associations, (ii) attention score (12.04-17.08%), derived from text mining of scientific literature, and (iii) heterogeneous graph walk (4.55-12.32%), which models relationships by exploring paths within a complex network of biological entities. These findings suggest that TargetPro effectively integrates biological signals derived from omics data with evidence extracted from human knowledge captured from publications, grant applications, and other relevant scientific literature. Such integration yields predictions that are more robust and reliable than those derived from any single data type alone, and generates more concrete hypotheses from target ranking, which is crucial for building confidence in the model among key stakeholders, including drug discovery scientists, potential licensing partners, and regulators.

The rapid growth of AI models for drug discovery has created an environment where tools are often evaluated using different datasets and performance metrics, making direct and fair comparisons difficult^35^. This lack of standardization slows scientific progress and complicates the selection of the most suitable tool for specific research needs. To address this gap, our framework introduces TargetBench 1.0 (**Figure 1**), a system designed for rigorous, reproducible, and transparent evaluation of target identification models. TargetBench 1.0 assesses models using criteria directly relevant to real-world drug development, such as the ability to retrieve known clinical targets or the drug development potential of newly proposed targets.

A key contribution of this work is the robust comparison of our specialized TargetPro model against a range of LLMs, including general-purpose models such as GPT-4o (See Related Links), Grok3 (See Related Links), and the domain-specific BioGPT^36^. This benchmarking, conducted under conditions designed to reflect real-world applications, provides critical insights into the relative performance and applicability of various target discovery models (refer to **Materials and Methods** for details). TargetPro achieved a top K clinical target retrieval of nearly 70% across all disease areas, significantly outperforming all tested LLMs (ranging 8-40%), and this advantage was maintained regardless of the number of targets requested (**Figure 5**). The decline in LLM precision with longer requested target lists reflects a typical behavior of generative models, which tend to be more accurate when prompted for a few top candidates but become less reliable when prompted to generate extensive lists that may include less certain predictions. This performance difference, however, is not a critique of LLMs, which have demonstrated remarkable capabilities in reasoning and scientific problem-solving^37^. Instead, it rather underscores a fundamental difference in model design and training. TargetPro is a supervised learning model explicitly trained to integrate multiple data modalities for a defined prediction task. In contrast, general-purpose LLMs are primarily pre-trained on vast volumes of unstructured text and are not inherently designed to discover informative weights for the specific categories of inputs used in this study without significant, specialized fine-tuning^12,38^. This underscores the importance of selecting the right tool for a given task and emphasizes the continuing value of specialized models for data-intensive scientific applications.

The ultimate goal of target identification is not just to rediscover known targets but to uncover novel candidates. The AI-derived score generated by TargetPro serves as a powerful initial filter, enriching the pool of potential targets for those with a higher intrinsic probability of progressing through clinical development (**Figure 5**). However, narrowing a broad list of candidates to a focused set of actionable targets requires incorporating additional metrics of ‘development potential’. These include assessment of druggability and supporting experimental evidence, both of which are critical for reducing failures in later stages of development^39^. Importantly, a target can be biologically relevant yet remain challenging to modulate with available technologies.

The analysis presented in **Figure 6** evaluates the development potential of the models using several criteria that are critical for generating ‘actionable intelligence’. While TargetPro consistently outperforms other models across most metrics, these metrics provide more than a simple comparison of performance between models and disease groups. They also reveal distinct attributes of ‘good’ novel targets across different therapeutic areas. The relative importance of different types of evidence varies by disease context, suggesting that target validation strategies should be context-dependent and disease-specific. For example, in a well-established field such as oncology, high-quality targets tend to have strong biological relevance and numerous available bioassays that enable rapid experimental validation (**Figures 6D and E**)^40,41^. In contrast, for high-attrition fields such as neurological and metabolic diseases, our analysis highlights drug repurposing potential as a particularly important de-risking strategy (**Figure 6C**)^42^.

To enable deployment of such evaluations across the drug discovery field, we have made the TargetBench 1.0 publicly accessible at https://www.targetbench.org for independent model assessment. This platform enables researchers to upload a ranked target list generated by their own proprietary models or discovery strategies. It then calculates the same key performance metrics used in this study (**Figures 5 and 6**) and presents the results graphically (**Supplementary Figure 2**). The user’s results are displayed alongside the benchmarked performance of the models reported in this study, enabling a direct comparison to established baselines.

In addition to benchmarking, our unified framework generates a ‘High Confidence Target List’ for each of the 38 diseases in the database (**Supplementary Table 2**). These lists are prioritized according to the AI score produced by the TargetPro model and validated through the TargetBench 1.0 system, providing actionable intelligence to guide drug discovery teams in selecting targets for costly and time-intensive experimental validation pipelines^30,43^.

While this framework represents a significant step forward, it is important to acknowledge its limitations. The analysis was performed on 38 diseases and does not include all human pathologies. Periodic updates to TargetBench 1.0 could enhance applicability to new indications. The definition of a ‘successful’ target was derived from historical clinical trial data, which may carry inherent biases and does capture cases where targets failed due to commercial rather than scientific factors. The retrospective nature of the dataset limits its ability to predict future performance, making it necessary for TargetBench 1.0 to incorporate alternative metrics, such as predicted druggability, the availability of approved drugs for other indications, and the number of bioassays for evaluation. A ‘time capsule’ approach, in which the results of predictive modeling are stored until a future date for reassessment, offers a valuable strategy for evaluating different target identification models. By comparing predictions to eventual clinical outcomes, this method enables assessment of model accuracy and elucidation of factors contributing to prediction failures. Furthermore, the 22 selected omics and text models, though extensive, do not include every possible data type^12^. Incorporating additional data, such as epigenomics, metabolomics, and single-cell derived, may further improve predictive accuracy^44,45^.

The modular design of our framework provides a clear path for future development and highlights its broad applicability. The workflow we applied to build TargetPro is not limited to the current 22 omics and text models. It can be readily applied to other multidimensional data sources, such as the models from the Open Targets platform (the current target lists were based on default, non-optimized rankings), to generate similarly enhanced predictive models^9,46,47^. Furthermore, the fundamental approach of integrating multiple weak signals to predict a specific outcome is highly versatile. By changing the input features and the target variable, this workflow could be adapted to address other key challenges in drug development, such as predicting patient response to a given therapy, identifying biomarkers for clinical trial stratification, or even forecasting the likelihood of success for a drug in a specific phase of clinical development.

A potential future direction is to integrate this framework into a fully automated, closed-loop discovery system, or a ‘self-driving lab’ ^48^. In such a system, TargetPro could act as the ‘AI brain’ to identify and prioritize promising targets. The results from automated *in vitro* knockout or inhibition experiments would then be fed back into the model for retraining and improvement, creating a continuous cycle of hypothesis generation, testing, and learning. Our framework provides the two core software components essential for such a system: the AI model that prioritizes targets for testing and an evaluation module that assesses the results. Together, these components create a dynamic and expandable platform that lays the foundation for a more AI-guided and automated approach to drug discovery.

## Related Links

Verge Genomics: https://www.vergegenomics.com/approach

TargetMATCH: https://www.owkin.com/targetmatch

HGNC database: https://www.genenames.org/

MedChemExpress: https://www.medchemexpress.com

GPT-4o: https://openai.com/index/gpt-4o-system-card/

Grok3: https://x.ai/news/grok-3

## Acknowledgements

We thank Ms. Elizaveta Ekimova for her technical assistance with figure design.

## Supplementary Figures

**Supplementary Figure 1.**
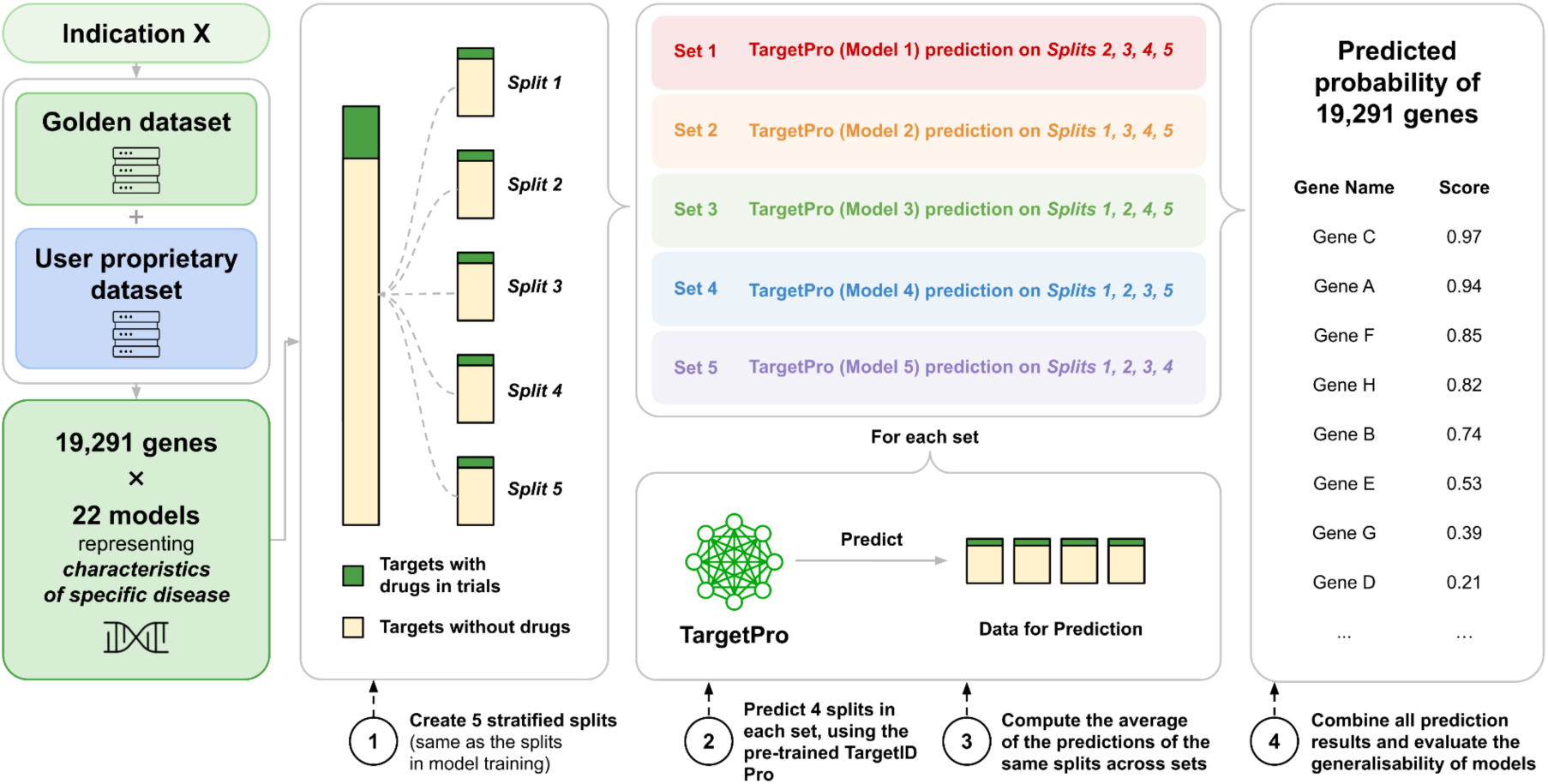
Workflow for generating and evaluating user case target lists. The schematic illustrates the process for generating target scores for a specific indication. First, the foundational ‘golden dataset’ is augmented with the ‘User proprietary dataset’ to create a combined dataset. This dataset is then processed to cover 19,291 genes, each described by 22 models, and is then divided into five stratified splits based on whether targets have associated drugs in clinical trials. Five sets of predictions are subsequently performed, where each model (TargetPro) predicts on the same set of genes it was trained on. The five sets of predictions are then combined to produce a final, comprehensive list of predicted probabilities for all 19,291 genes. Finally, these consolidated prediction results are used to evaluate the overall quality and performance of the model.

**Supplementary Figure 2.**
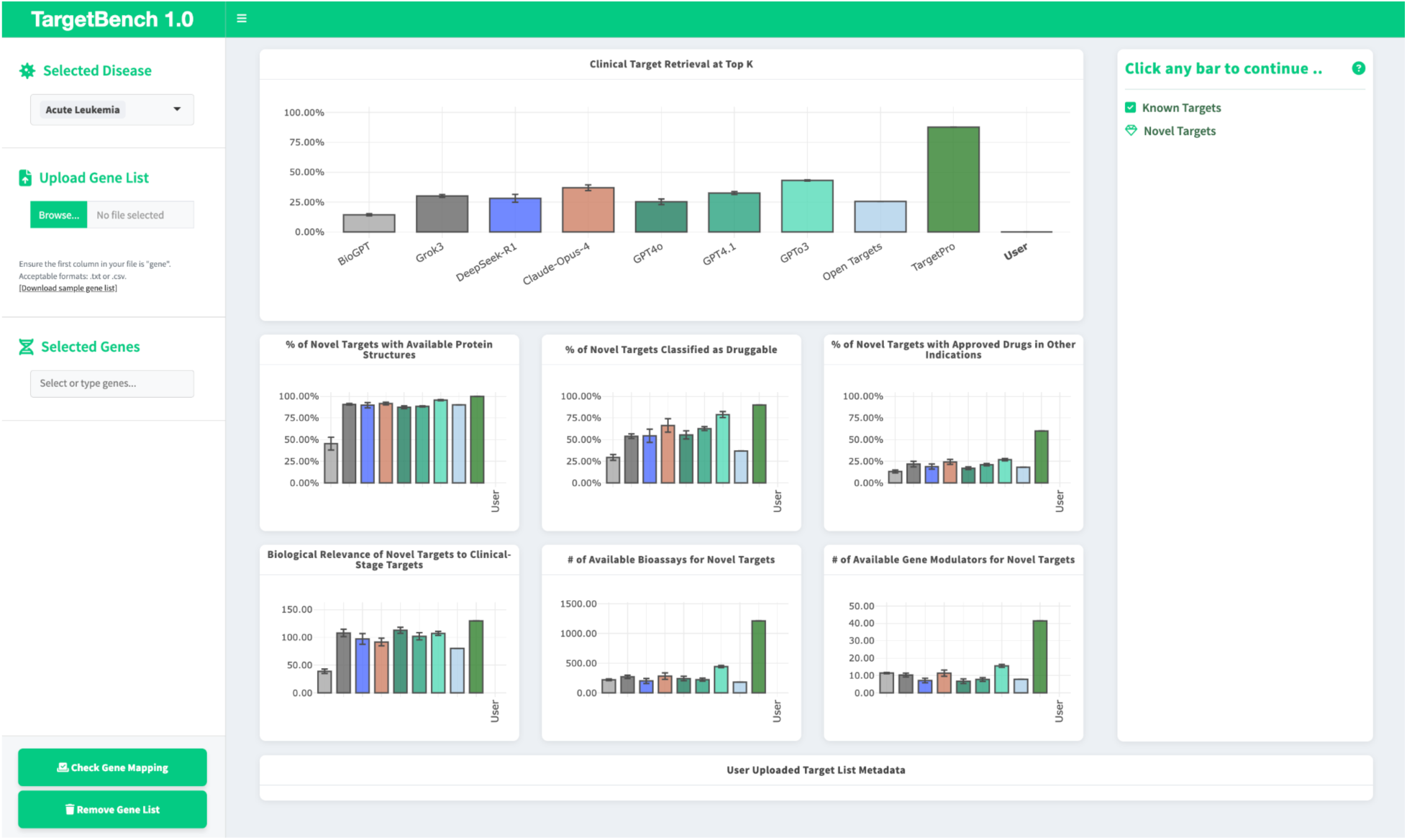
TargetBench 1.0 web UI

